# Proteome mapping of the human pancreatic islet microenvironment reveals endocrine-exocrine signaling sphere of influence

**DOI:** 10.1101/2022.11.21.517388

**Authors:** Sara JC Gosline, Marija Velickovic, James Pino, Le Z. Day, Isaac K. Attah, Adam C. Swensen, Vincent Danna, Karin D. Rodland, Jing Chen, Clayton E. Matthews, Martha Campbell-Thompson, Julia Laskin, Kristin Burnum-Johnson, Ying Zhu, Paul D. Piehowski

## Abstract

The need for a clinically accessible method with the ability to match protein activity within heterogeneous tissues is currently unmet by existing technologies. Our proteomics sample preparation platform, named microPOTS (Microdroplet Processing in One pot for Trace Samples), can be used to measure relative protein abundance in micron-scale samples alongside the spatial location of each measurement, thereby tying biologically interesting proteins and pathways to distinct regions. However, given the smaller sample number and amount of tissue measu red, standard mass spectrometric analysis pipelines have proven inadequate. Here we describe how existing computational approaches can be adapted to focus on the specific biological questions asked in spatial proteomics experiments. We apply this approach to present an unbiased characterization of the human islet microenvironment comprising the entire complex array of tissues involved while maintaining spatial information and the degree of the islet’s sphere of influence. We identify specific functional activity unique to the pancreatic islet cells and demonstrate how far their signature can be measured. Our results show that we can distinguish pancreatic islet cells from the neighboring exocrine tissue environment, recapitulate known biological functions of islet cells, and identify a spatial gradient in the expression of RNA processing proteins within the islet microenvironment.

## Introduction

The islets of Langerhans are endocrine micro-organs embedded within a mostly exocrine pancreas, comprising roughly two percent of the pancreas by mass. Islets have been studied for decades primarily because of their involvement in diseases such as diabetes and obesity. Until recently, in-depth protein profiling of pure islets has been very difficult, partly due to their diminutive size and limited compositional make up. Recent cutting-edge technologies have greatly enhanced our understanding of the islet proteome by isolating islets from their surrounding tissues allowing them to be studied down to near single-cell resolution^1–3^. In-depth proteomic studies of acinar cell tissues from the exocrine pancreas have also been demonstrated in the past^4,5^. However, despite their encapsulated nature, islets do not act entirely independently and rely on the surrounding exocrine microenvironment for feedback signaling and crosstalk^6,7^. A characterized islet-acinar portal system directly facilitates islet hormone dispersion in nearby acinar cells. For example, acinar cells are known to contain islet-hormone-specific receptors that regulate acinar function and under the right conditions saturate with locally high concentrations of insulin and somatostatin^8^. In addition to insulin and somatostatin, other humoral factors including pancreastatin and ghrelin, and several neurotransmitters (nitric oxide, peptide YY, substance P, and galanin) have been shown to be involved in this islet-acinar connection; regulating the functions of each tissue type^9^. Through causes we do not yet understand, people with type 1 diabetes and their first-degree relatives have also been shown to have overall reduced pancreatic volume compared to matched controls and some evidence supports exocrine pancreas atrophy and exocrine insufficiency in people with long term T1D^10–12^. The more we learn about this endo-crine-exocrine/islet-acinar connection the greater its importance appears to be in understanding diseases involving the pancreas. However, until now no unbiased spatially resolved method has been available for deep proteomic investigations of the islet microenvironment encompassing all the cell/tissue types involved this complex system.

Large bulk samples of pancreatic tissues quickly dilute and drown out the contribution of the islet signature. To gain deeper understanding of the biological signaling and interactions that underlies this endocrine-exocrine connection, it is critical to study the tissues with their original spatial context intact (i.e., in vivo)^13,14^. Currently, few technologies are available to study the heterogeneity of biological signaling across a tissue sample. Although there are several powerful techniques for measuring transcripts with high depth and spatial resolution, transcripts often don’t correlate well with protein expression^15^. Existing technologies for spatially resolved protein measurements mainly rely on the use of tagged antibodies, such as Immunohistochemistry^16^, CyTOF^17^ and CODEX^18,19^. While these technologies are highly effective and can provide single-cell level or better spatial resolution, protein coverage is limited by the availability of reliable antibodies and the multiplexing limit of the labels. Imaging mass spectrometry (MALDI, Laser Ablation) is also a powerful tool for protein mapping that does not depend on antibody recognition; but due to the direct coupling to the mass spectrometer, these techniques are limited in their dynamic range and accuracy of quantitation^20–24^.

Over the last decade, improvements in sensitivity and sample handling for LC-MS proteomics have enabled spatially resolved measurements^25–31^ and have extended to more difficult to analyze samples such as formalin fixed paraffin embedded (FFPE) tissue^26,32,33^. These approaches are particularly attractive as they offer a comprehensive, quantitative protein profile without *a priori* knowledge of the proteins of interest. In our lab, we have successfully combined laser capture microdissection and the nanoPOTS approach^34,35^, to enable in-depth proteome imaging. This initial effort resulted in a platform capable of quantifying >2000 proteins at 100 μm spatial resolution without the use of antibodies or labels^36^. To further improve the depth of protein coverage, we next incorporated tandem mass tags (TMT) and nanoflow fractionation and concatenation (nanoFAC^37^) to our proteome imaging workflow. To collect and process enough protein material to facilitate robust fractionation, we scaled up the platform to the microliter scale^38,39^, and incorporated a TMT carrier channel^40,41^ into the image plexes. These changes increased protein coverage to >5000 proteins while maintaining high-quality quantitative information, thus enabling spatially resolved, unbiased interrogation of biological signaling.

Robust computational analysis tools for the analysis of data from these evolving technologies are less established. Technologies such as CyTOF and CODEX have proprietary software packages that are sold with their technology^19^, though there are many open source tools that leverage these data for the purposes of characterizing cells by their protein expression via flow cytometry^42–44^. There are also computational packages designed for spatially resolved transcriptomics data that can be leveraged for proteomic analysis, including those that enable mapping and analysis of imaging data^45,46^ as well as those that link the two data modalities to improve protein identification and clustering^47^. Existing tools, however, are limited to the study of pre-formatted image data based on the established platform (e.g. Visium^48^) and therefore are not easily applied to microPOTS data.

In this work, we demonstrate that the potential utility for microPOTS spatial proteomics to be employed in a clinical setting through the study of multiple pancreas regions within a single patient. We enhance existing computational tools to show how these measurements can be used to robustly measure activity across disparate regions within a single pancreas, and to make biologically functional hypotheses from these data. In addition to showing the increased insulin signaling activity we know to be present within the islet cells, we also identify specific immune related processes and RNA processing activities that could not be captured at the transcriptomic level and can be studied further in disease settings such as pancreatic cancer or diabetes.

### Experimental Procedures Experimental Design

Our overall analysis pipeline is depicted in **Figure 1** and described below. Seven proteome images were created from seven different regions of a pancreas tissue section taken from a healthy human donor. Images consist of 9 tissue “voxels” created by dissecting a 3 x 3 grid from the tissue collected directly into corresponding wells in a microPOTS chip (**Supplemental Figure 1**). Imaging areas were created from regions containing a singular group of islet cells to interrogate the islets and their unique microenvironment. Grids were sized to capture the islet within a single voxel consisting almost entirely of islet cells. Below we describe how we captured the data to identify islet-specific signaling activity.

**Figure 1.**
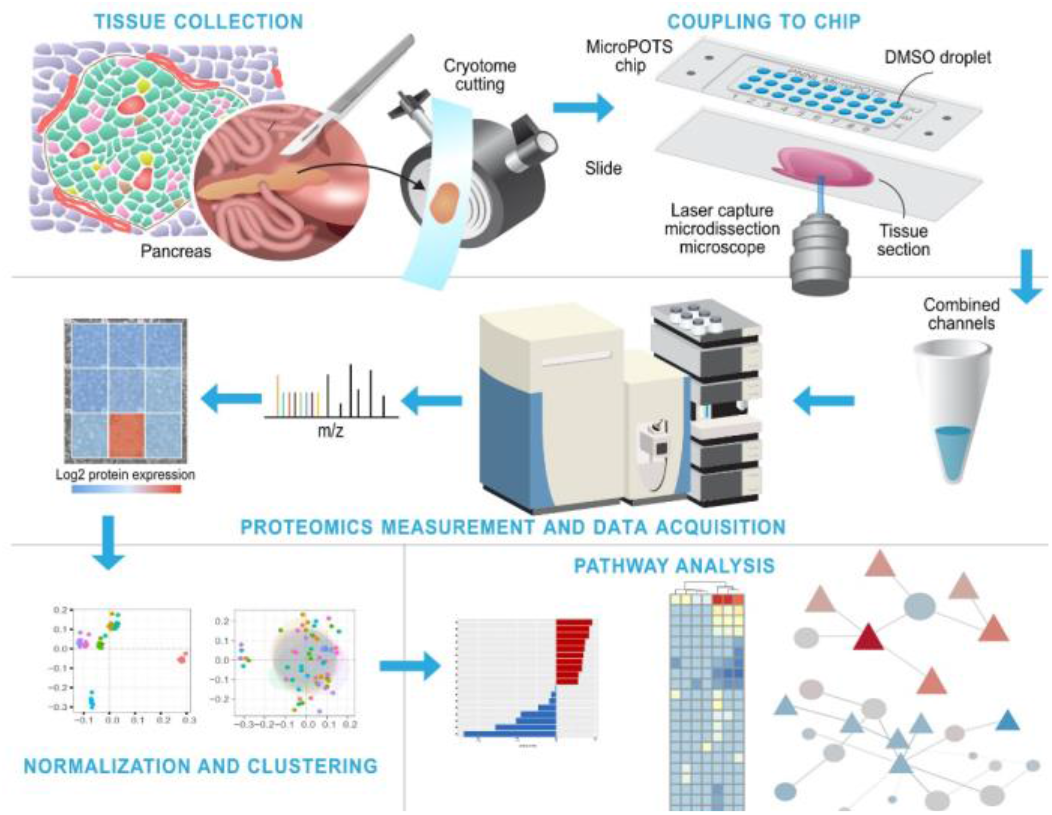
Overview of our experimental procedures. Top: tissue collection and coupling to chip requires laser capture microdissection of flash frozen pancreatic samples enables dissection of each ‘voxel, into each microwell for sample preparation. Middle: each well is individually prepared for TMT labeling and MS/MS. To measure relative protein abundance for each voxel. Bottom: individual voxels are annotated to carry out pathway enrichment and network analysis.

### Tissue collection and coupling to chip

#### Tissue collection

Samples were washed with the gradient of ethanol solutions (70%, 96%, and 100% ethanol, respectively) to dehydrate the tissue sections and to remove embedding material. Human pancreas tissue for microPOTS imaging was obtained from a 17-year-old male donor. The donor was selected based on our eligibility criteria established by the HuBMAP consortium^49^. Organ recovery and tissue processing were performed at University of Florida per standard protocol^50^. Briefly, pancreas was sliced into 0.5 cm-thick tissue segments, subdivided, and immediately frozen in Carboxymethylcellulose (CMC, prepared in Cryotray molds that were prechilled on dry ice/isopentane slurry^51^. Frozen CMC tissue blocks were stored at −80°C until sectioning. CMC embedded human pancreas tissue was cut to 10-μm-thick slices using a cryostat and collected on PEN membrane slides. Serial sections were shipped on dry ice to PNNL for further microPOTS imaging.

#### Laser capture microdissection (LCM)

Sample dissection and “voxel” collection were completed using a PALM Microbeam system (Carl Zeiss Microimaging, Munich, Germany) which contains a RoboStage for high-precision laser micromanipulation in micrometer range and a PALM RoboMover that collects voxel samples directly into the wells of the microPOTS chip. Microwells were preloaded with 3 μl of dimethyl sulfoxide (DMSO) that served as a capturing medium for excised voxels.

For proteomics imaging experiments, we first stained a 10 μm thick adjacent human pancreas section using Periodic Acid-Schiff (PAS) staining kit following the manufacturer’s protocol. The staining for the confident determination of islet and acinar tissue regions when observed using bright-field microscopy. Informed by the islet localization from the serial PAS-stained section, a 3×3 grid was created over an islet and the surrounding acinar tissue. The grid was arranged to capture the whole islet in a single pixel of approximately 200 μm x 300 μm dimensions while the surrounding 8 pixels contained exclusively acinar tissue. Voxels were dissected using the grid mode and collected directly into corresponding microwells of the chip. A carrier sample was also collected that contained a similar-sized islet and surrounding acinar tissue with a total area equivalent to the entire grid size, which was ~500,000 μm^2^.

#### Proteomics sample processing in a microdroplet

All sample handling steps, from extraction through to TMT labeling, were carried out on-chip by manual pipetting. Evaporation during preparation was minimized by cooling during dispensing of reagents, using a humidified chamber for incubation steps, and sealing the chip with a contactless cover and wrapping in aluminum foil. Tissue voxels were incubated at 75°C for 1 hr to remove DMSO solvent. Next, 2 μl of extraction buffer containing 0.1% DDM, 0.5×PBS, 38 mM TEAB, and 1 mM TCEP was dispensed to each well of the chip, followed by incubation at 75°C for 1 hr. We then added 0.5 μl of 10 mM IAA solution in 100 mM TEAB to reach a final concentration of 2 mM IAA followed by incubation at room temperature for 30 min. Samples were subsequently digested by dispensing 0.5 μl of an enzyme mixture (10 ng of Lys-C and 40 ng of trypsin in 100 mM TEAB) and incubated at 37 °C for 10 h. TMT-11 plex reagents were re-suspended in anhydrous acetonitrile at a concentration of 6.4 μg/ μL. 1 μl of each TMT tag was used to label voxel samples. Following our experimental design, each plex/image was created by leaving the 130N channel empty and using the 131N channel for the carrier sample, 128N channel was used for the islet voxel and the other 8 channels for the acinar tissue voxels. The peptide-TMT mixtures were incubated for 1 h at room temperature, and the labeling reaction was quenched by adding 1μl of 5% HA in 100mM TEAB and incubating 15 min at room temperature. All samples were then pooled together, brought up to the final 1% FA, then centrifuged at 10,000 rpm for 5 min at 25 °C. Finally, the pooled sample was transferred to an autosampler vial, and dried in a speed vac.

#### Reagents and Chemicals

Microwell chips with 2.2 mm well diameter were manufactured on polypropylene substrates by Protolabs (Maple Plain, MN). LC-MS grade water, formic acid (FA), iodoacetamide (IAA), Triethylammonium bicarbonate (TEAB), TMT-10plex and TMT11-131C reagents, Anhydrous acetonitrile, Tris(2-carboxyethyl)phosphine hydrochloride (TCEP-HCl), and 50% Hydroxylamine (HA) were all purchased from Thermo Fisher Scientific (Waltham, MA). N-Dodecyl β-d-maltose (DDM), DMSO (HPLC grade), and Phosphate-Buffered Saline (PBS) and PAS staining kit were purchased from Sigma-Aldrich (St. Louis, MO). Both Lys-C and trypsin were purchased from Promega (Madison, WI). Ethanol was purchased from Decon Labs, Inc (King of Prussia, PA).

### Proteomic measurement and data acquisition

Nanoflow LC-fractionation. Prior to injection, samples were resuspended in 62 μl of 0.1% formic acid. High pH fractionation was performed off-line by loading 50 μl of the sample onto a precolumn (150 μm i.d., 5 cm length) using 0.1% formic acid at a flow rate of 9 μL/min for 9 min. The sample is then pushed onto the LC column (75 μm i.d., 60-cm length) using the separation gradient. Precolumn and column were packed inhouse with 5-μm and 3-μm Jupiter C18 packing material (300-Å pore size) (Phenomenex, Terrence, USA), respectively. An Ultimate 3000 RSLCnano system (Thermo Scientific) was used to deliver gradient flow to the LC column at a nanoflow rate of 300 nl/min. 10 mM ammonium formate (pH 9.5) was used as mobile phase A and acetonitrile as mobile phase B. Eluted fractions were collected using a HTX PAL collect system into autosampler vials preloaded with 25 μl 0.1% formic acid and 0.01% (m/v) DDM. The PAL autosampler allows concatenation on-the-fly by robotically moving the dispensing capillary among 12 collection vials. A total of 96 fractions were concatenated into 12 fractions. Vials were stored at −20 °C until the following low-pH LC-MS/MS analysis.

#### LC-MS/MS peptide analyses

LC-MS/MS analysis was carried out using the Ultimate 3000 RSLCnano system (Thermo Scientific), coupled to a Q Exactive HF-X (Thermo Scientific) mass spectrometer. Full MS1 scans were acquired across scan range of 300 to 1,800 m/z at a resolution of 60,000, combined with a maximum injection time (IT) of 20 ms and automatic gain control (AGC) target value of 3e6. Data dependent MS2 scans were collected using a top 12 method with a resolving power of 45,000, a maximum injection time of 100 ms, and AGC target value of 1e5, with the isolation window was set to 0.7 m/ z and dynamic exclusion time was set to 45 s to reduce repeated selection of precursor ions.

#### Data Analysis

InstrumentRAW files were first processed using MSConvert to correct mass errors^52^. Corrected dspectra were searched with MS-GF + v9 88 1 ^53,54^ against the Uniprot human database downloaded in March of 2021 (20,371 proteins) and a list of common contaminants (e.g., trypsin, keratin). Partially tryptic search setting was used and a ± 20 parts per million (ppm) parent ion mass tolerance. A reversed sequence decoy database approach was used to control the false discovery rate. Carbamidomethylation (+57.0215 Da) on Cys residues, and TMT modification (+229.1629 Da) on N terminus and Lys residues were considered as static modifications. Oxidation (+15.9949 Da) of Met residues was set as a dynamic modification. Identifications were first filtered to a 1% false discovery rate (FDR) at the unique peptide level, and a sequence coverage minimum of 6 per 1000 amino acids was used to maintain a 1% FDR at the protein level after assembly by parsimonious inference.

TMT 11 reporter ions area under the curve (AUC) intensities were extracted using MASIC software^55^. Extracted intensities were linked to peptide-to-spectrum matches (PSMs) passing the FDR thresholds described above. Relative protein abundance was then calculated as the ratio of sample abundance to the median abundance of the protein across all datasets, using the summed reporter ion intensities from peptides that could be uniquely mapped to a protein. While we typically transform/zero-center the relative abundances for each gene relative to the final abundance value, we asked in the next section if that was necessary here.

### Normalization and statistical rationale

We compared the batch effects and made normalization decisions via PCA and clustering. Each image was annotated by its sample number and position in the grid – each element of the grid was labeled as ‘islet’ if it contained islet cells, ‘proximal’ if the voxel was adjacent to the islet, and ‘distal’ otherwise. The first two principal components were used to assess batch effects in **Figures 2B–C** and **2E–F.**

**Figure 2.**
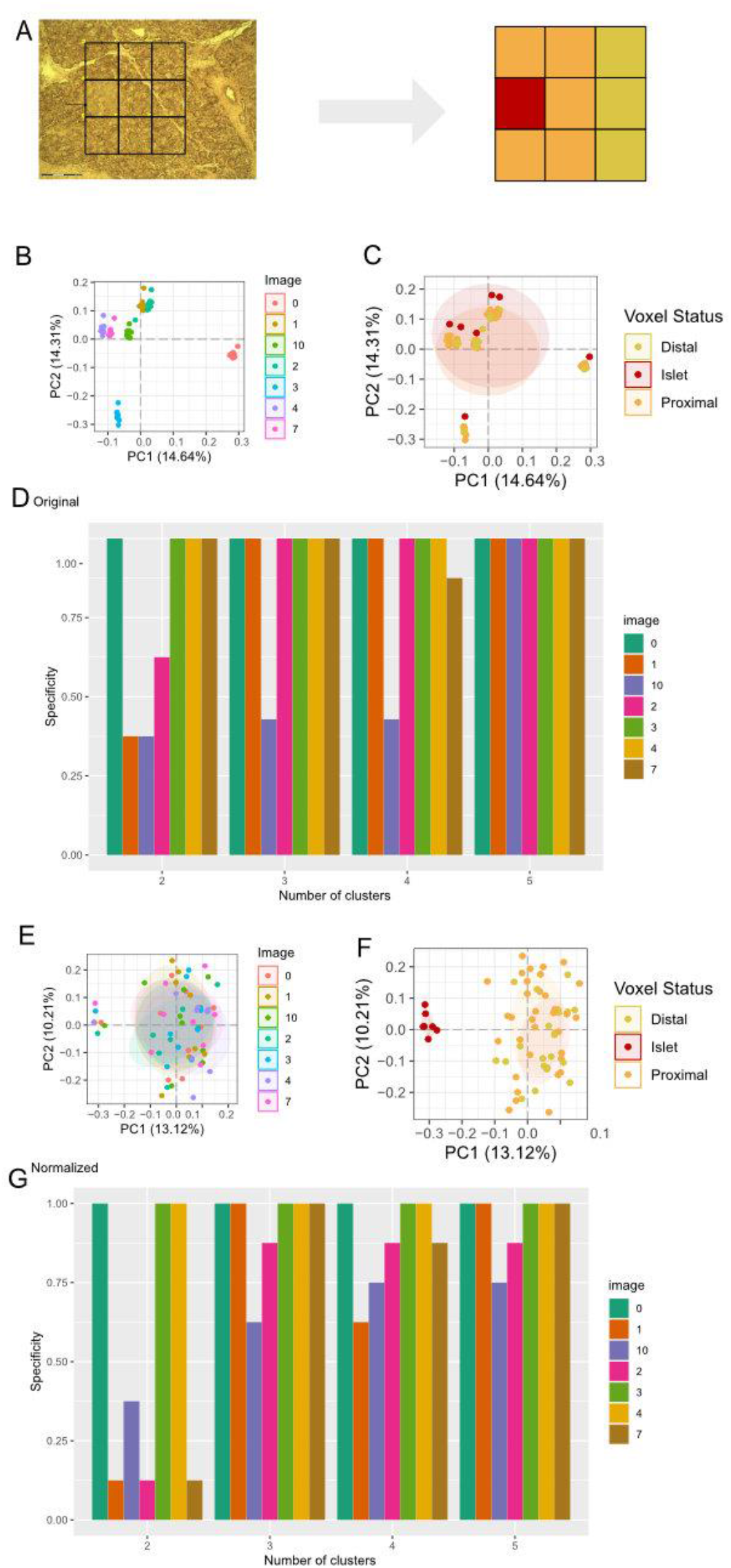
Statistical analysis. (A) stained pancreatic region and 3×3 grid placement centered around islet cell, indicated by arrow. Scale bar is 150 uM. (B) Principal component analysis (PCA) of uncentered voxels shows grouping by each image compared to (C) the islet annotation. (D) BayesSpace specificity (y-axis) colored by image number, where x-axis represents number of clusters used in the algorithm. (E-F) shows PCA of median centered images clustering by islet cell annotation rather than image. (G) shows BayesSpace specificity.

The BayesSpace^47^ clustering algorithm was used to cluster the voxels into 2, 3, 4, and 5 clusters based on position in the grid and proteomics measurements. To compare median centering of the individual voxels, we measured specificity of each clustering approach. If the islet-containing voxel was the only one in its cluster, the specificity was 1. Otherwise, it was lower. These results are depicted in **Figures 2D** and **2G.**

### Pathway analysis

To compute differences between the islet cell-containing voxel and other regions, we used the limma package with pooled images (with the annotation described as above). This enabled each image to be its own biological replicate, particularly of the islet cells, which were only measured in one voxel per image. We then compared analysis of regions using the upsetR^56^ tool to compare expression differences across annotated regions, then employed leapR^57^ pathway enrichment tool to identify specific pathways that were up regulated in islets across all 7 images using the enrichment_in_sets parameter with the KEGG, Reactome, and GO Biological Process pathways. We used the same pathways to do the enrichment_in_order analysis in our variancebased and distance-based analysis. The network analysis leveraged the PCSF R tool^58^ based on the approach described previously^59^.

## Results

Here we describe the analysis pipeline by which we investigate the proteomics imaging of the microPOTS framework to enable interpretation in biological use cases.

### Spatial proteomics across multiple images enables quantitation of proteins at high resolution

We collected seven distinct samples from a single human pancreas and disected each sample into a 3×3 grid for 9 “voxels” for each image (**Supplemental Figure 1**). Images were stained with Periodic Acid-Schiff (PAS) to identify the islet (**Figure 2A**) and an adjacent sample was captured for proteomics. After running each of the 9 samples for the 7 images through our mass spectrometry pipeline (see Methods), we captured ~6000 distinct proteins (**Supplemental Table 1**).

We compared the clustering of the individual voxels as described above and determined that, despite the batch effect observed between images, we could still robustly identify the islet containing voxels (**Figure 2**). In fact, using the BayesSpace clustering algorithm, we were able to identify islets with more specificity without normalization (**Figure 2C** and **2E**). As such, we decided to keep the original log2 ratio values.

We first sought to identify protein targets we knew have high cell-type specificity. **Figure 3A** shows the uncentered log2 fold change values of Insulin, a hormone released in islet cells across each of the 7 images, while **Figure 3B** shows the expression of glucagon. Each islet cell is annotated with a solid box as derived from the imaging data. As expected, both insulin and glucagon are highest in the islet cells across of the 7 images. Note that we were unable to resolve beta cells (insulin) from alpha cells (glucagon) at this level of spatial resolution, and that insulin is also highly expressed in a non-islet-containing voxel in image 3, likely due to an islet in another region that was not measured.

**Figure 3.**
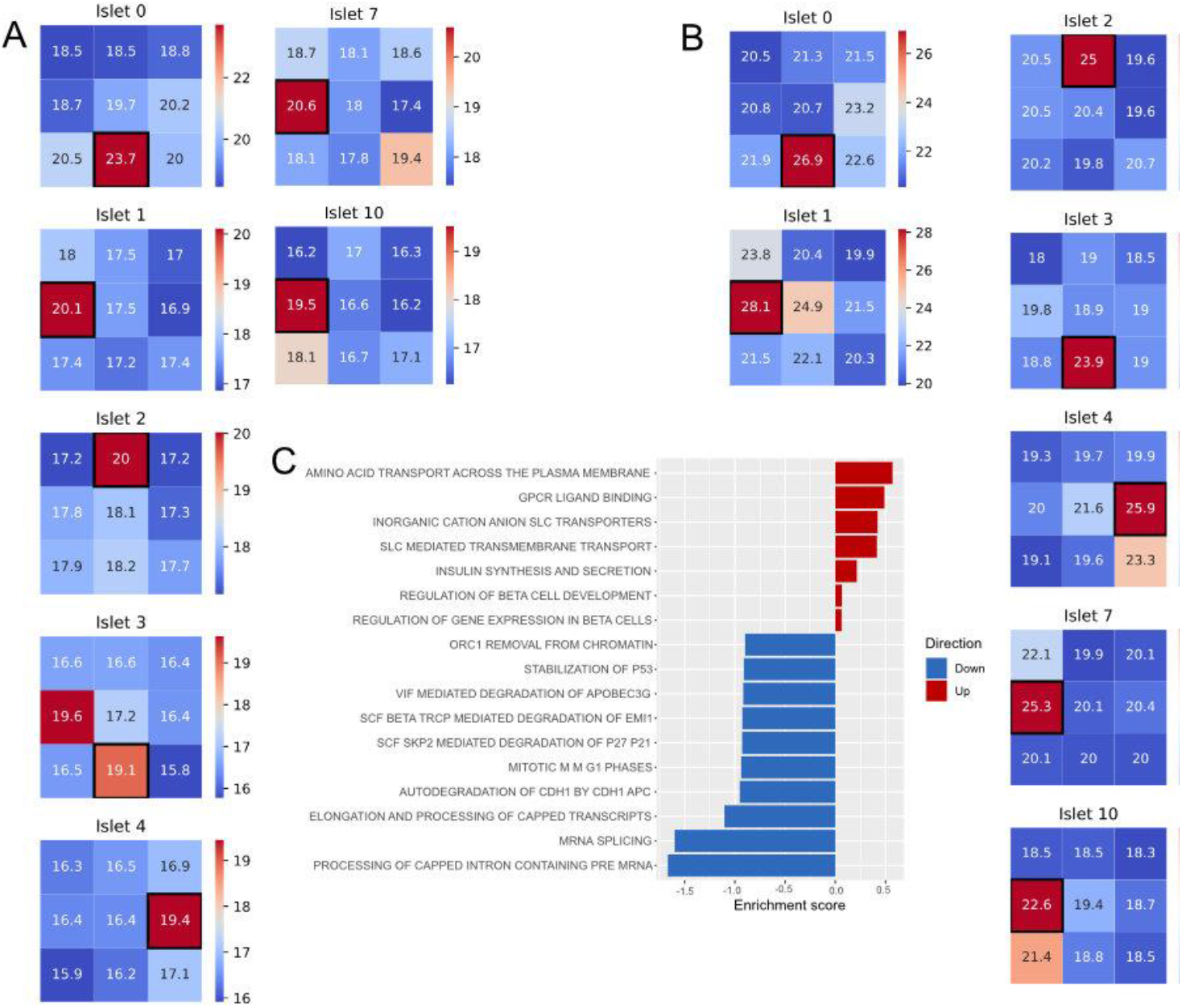
Mapping protein expression to islet regions. (A) Expression of Insulin protein across all 7 images, with islet highlighted by black box. (B) Expression of Glucagon across all 7 images, with islet region highlighted in black. (C) Gene set enrichment of biological pathways of proteins ranked by variance across all 63 voxels.

Lastly we assess the variance of each protein across all samples to determine if proteins in some pathways are changing more than others. We then used ranked gene set enrichment (see Experimental Procedures) to identify if the variance of proteins corresponded to some pathways. The results, depicted in **Figure 3C** show that most of the pathways that are highly variable – depicted in red – are related to insulin or beta cells, suggesting that the primary differences in protein expression are between the beta cells and acinar tissue.

### Imaging pooling enables capture of islet-specific enrichment patterns

The protein expression signature of islets is substantially different than the neighboring acinar tissue. We determined the most significant pathways that differed, either increased or decreased expression in islets compared to the neighboring microenvironment. The voxel grids were positioned so that the majority of one of the central edge voxels encapsulated a complete islet. The remaining voxels were annotated based on their spatial distance from the islet voxel. For each image we annotated the five voxels immediately surrounding the islet as ‘proximal’ and the remaining three voxels as ‘distal’ (**Figure 2A**). We then grouped the expression of all voxels by these three annotations. We examined differences in the non-islet regions, as shown in the upset plot in **Figure 4A**. As expected, we found the majority of differentially expressed proteins between the islet and other cells were also differentially expressed between the islets and the proximal/distal regions when compared independently.

**Figure 4.**
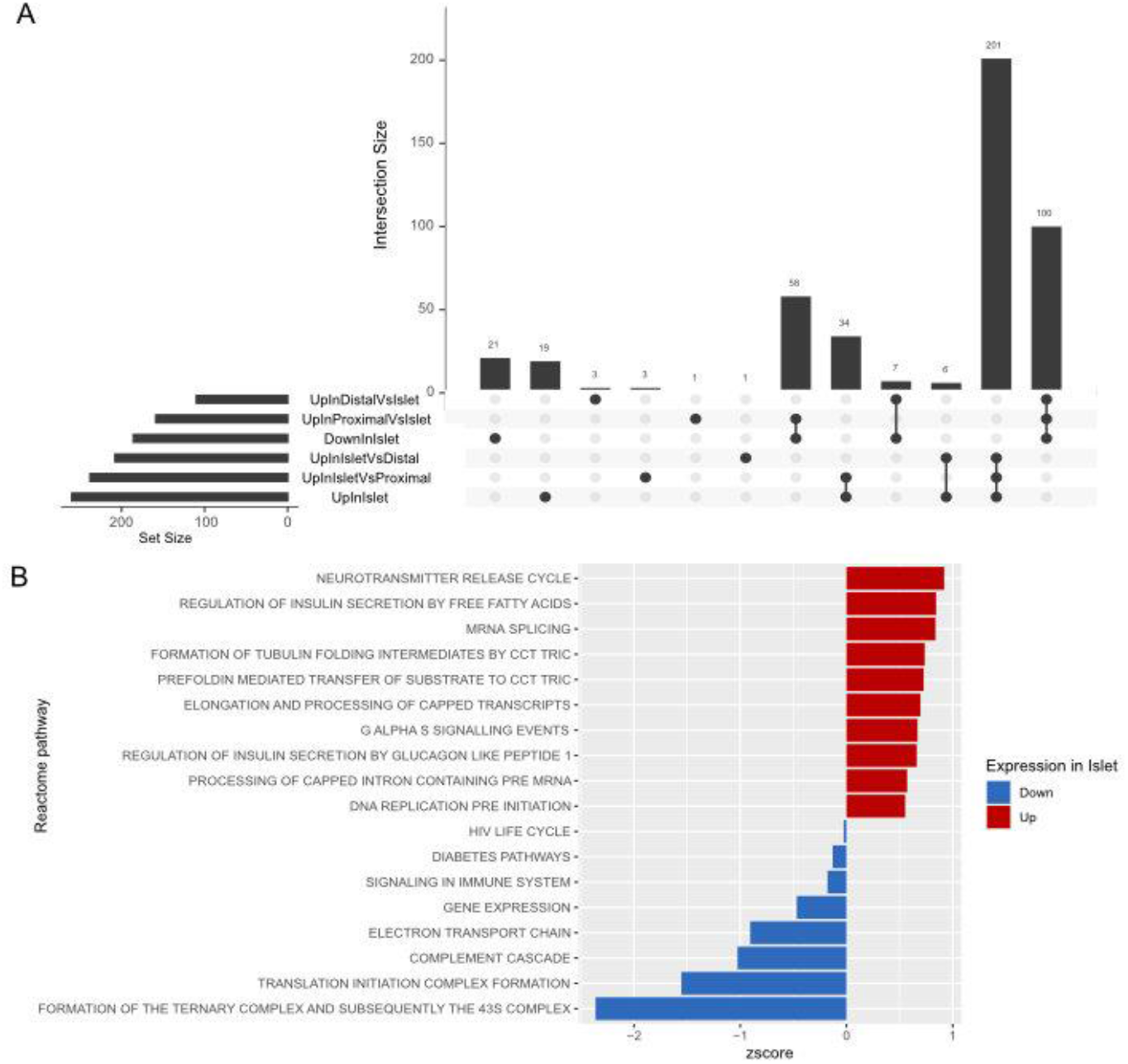
Image-pooling differential expression analysis. (A) shows pairwise differential expression between each region. (B) Pathways enriched in the islet cells compared to the union of distal and proximal regions.

We then evaluated the pathways that were up-regulated in islet cells compared to the non-islet cells. The results, shown for Reactome pathways in **Figure 4B** depict the most statistically significant terms that were up-regulated in the islets (red) and down-regulated (blue). Confirming the role of the islet cell in insulin signaling^60^, we see regulation of insulin secretion as one of the most enriched pathways in the islet cells. We also observed enrichment in RNA splicing^61,62^ and pre mRNA capping as up-regulated. In the acinar tissue, we see up-regulation of immune signaling and the complement cascade, suggesting a role in immune machinery in the pancreas.

### Network analysis implicates related proteins in key islet pathways

Given the number of proteins derived from the microPOTs measurements (~6000 proteins per voxel) we explored network inference tools to determine if we could infer biological signaling pathways based on the protein expression alone. Specifically, we used the Prize-collecting Steiner tree algorithm (see Experimental Procedures) to identify the network implicated by proteins up-regulated in islets (red, **Figure 5A**) and down-regulated in islets (blue, **Figure 4B**). As input to the algorithm, we had 261 up-regulated proteins and 184 down-regulated proteins. The resulting networks were 216 and 179 nodes, respectively, as the algorithm removed proteins that were not connected to others in the interactome and added proteins that maximized the connection of differentially expressed proteins in the network.

**Figure 5.**
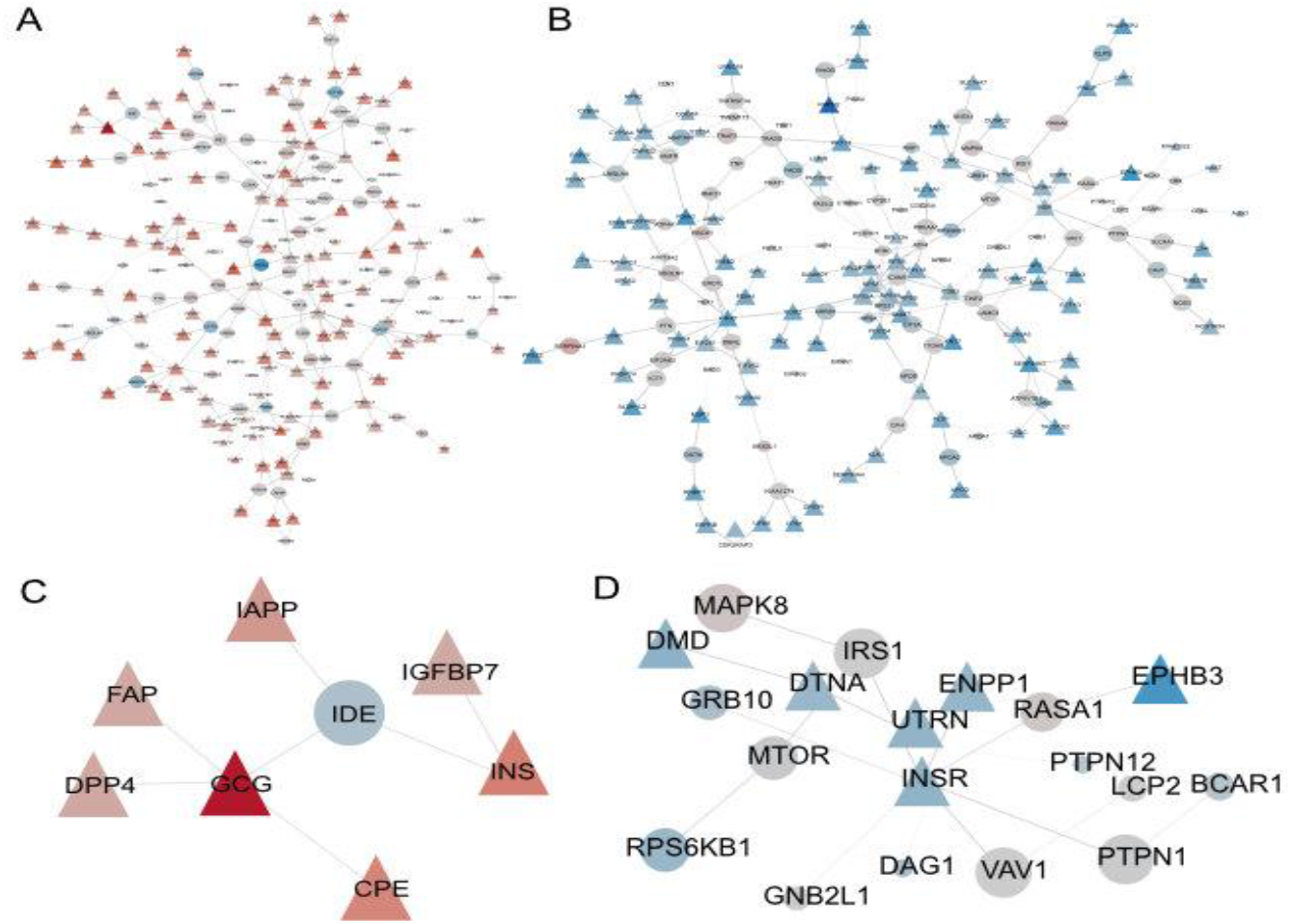
Network analysis. (A) Protein interaction network of 216 nodes uniquely active islet cells across images. Red indicates degree of up-regulation in islets, blue indicates down-regulation. Triangles indicate protein is differentially expressed, while circles mean the protein is inferred to be active. (B) Protein interaction network of 179 proteins that are down-regulated in islets. Colors and shapes as in (A). (C) Subnetwork of (A) highlighting the role of insulin (INS) and glucagon (GCG). (D) subnetwork of (B) highlighting the insulin receptor (INSR).

This approach allows us to investigate specific nodes implicated (i.e., not detected experimentally but added via the algorithm) in the network (circles in **Figure 5**). Specifically, in the islet network, **Figure 5C**, we found IDE, an Insulin-degrading enzyme. This protein is implicated in the network as an interactor of many highly expressed proteins including Insulin, Glucagon, and Islet amyloid peptide. Despite being a clear regulator of these proteins^60^ Insulin-degrading enzyme, while somewhat downregulated (but not statistically so) in the islets, is essential for the regulation of these proteins and therefore belongs in the signaling network. Similarly, we found key proteins that interact with the insulin receptor, which is up-regulated in the acinar tissue (**Figure 5D**). Here we found this protein to be centrally interacting with PTPN1, VAV1, and RASA1, all proteins whose change in expression was not statistically significant, and without VAV1 even being detected despite it being shown to support Beta cell maturation^63^.

### Distance based metric reveals distinct RNA processing changes as distance from islet increases

To further exploit the spatial relationship between voxels, we searched for signals that permeated through the tissue from the voxel. To do this, measured the spearman rank correlation of the expression of each protein to the voxels distance to the islet cell. We then searched for biological pathways using the Gene Ontology biological processes (**Figure 6A**) or the KEGG pathways (**Figure 6B**) that were enriched in proteins with a high positive correlation (more active farther from islet) or a highly negative correlation (more active closer to islet). We computed the results for each of the seven images, to ensure that we were getting similar results, shown in **Figure 6**. We found numerous pathways, such as rRNA processing/metabolism and the ribosome, to be enriched across all 7 images, which suggests that these proteins are more highly expressed the farther the voxel is from the islet cell.

**Figure 5.**
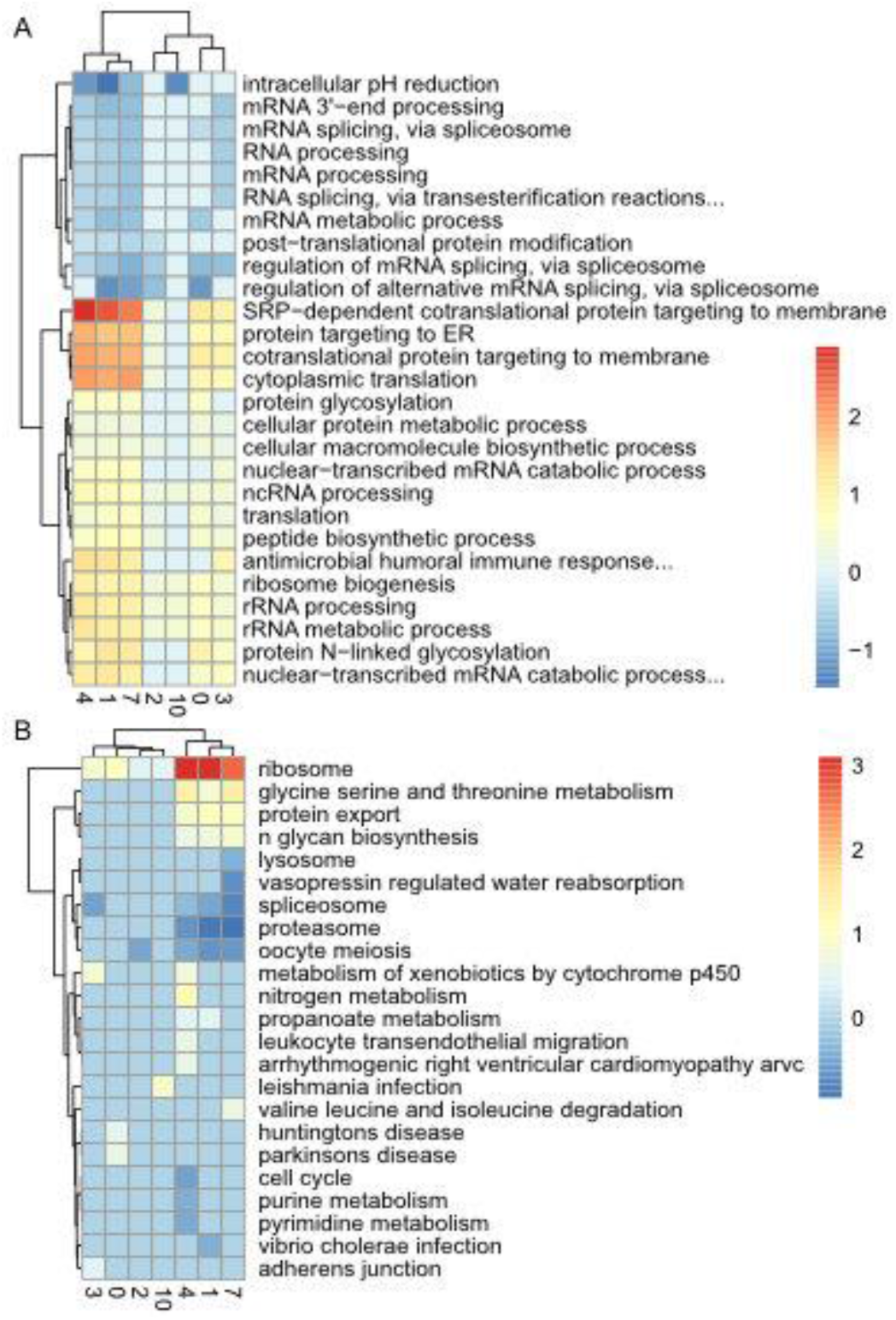
Distance-based analysis. For each image, we computed the correlation statistic between the protein of interest and the Manhattan distance to the islet voxel. We then ranked the proteins by correlation and computed gene set enrichment on (A) GO biological process and (B) the KEGG pathways. Red indicates positive correlation with distance, enriched farther from islet, while blue indicates negative correlation, or enrichment closer to islet

When compared to the baseline functional enrichment analysis (**Figure 4**) the distance-based approach shows the same up-regulation of RNA splicing but identifies specific elements of the splicing activity within the ribosome. Additionally, it identifies other RNA processes such as rRNA processing, ER protein targeting, and ncRNA processing, as being up-regulated farther from the islet cells. This clarifies the recent findings of RNA processing regulation changes in pancreatic beta cells^64^.

## Discussion

Here we introduce how the microPOTS spatial proteomics platform can be utilized in a clinical setting to characterize specific biological pathways that are uniquely expressed in pancreatic islet cells. We can robustly characterize ~6000 proteins in each sample and identify, within each sample, islet-specific biological processes. Our computational analysis enhances initial proteomic resolution through network integration and distance analysis.

These results highlight the distinction between insulin secretion, an exclusively islet cell activity, and insulin signaling, which is clearly enriched in the neighboring acinar cells. Network analysis highlights PTPN11, VAV1, and RASA1 as key hub proteins which influence the down-stream response to upstream stimuli^63^; for example, these hub proteins could act to funnel contrasting signals from insulin and glucagon to the same set of distal cellular responses, providing a mechanism for opposing physiological consequences of these effectors. Interestingly, the observed increase in foundational RNA processing activities with increasing distance from the islets confirms recent findings of ribosomal changes^64^ and suggests that exocrine cells at a distance from islets are predominantly engaged in normal activities associated with cell growth and renewal, while acinar cells in closer proximity to islets may be more specialized for signal transduction. Clearly these are hypothesis generating observations, and substantiation of these hypotheses would require careful mechanistic experiments, possibly using a spatially controlled system such as pan-creas-on-a-chip systems. The value of this unbiased spatial proteomic approach is that it suggests targets for genetic manipulation in future experiments.

In summary, we believe the technology and analysis procedures described herein enable a diverse set of applications of proteomics in the clinical setting. There are many diseases, such as cancer or Type 1 diabetes, in which a small group of cells can cause a large amount of damage. As such, these technologies are imperative to enable the study of specific signaling activities that enable these cells to affect the neighboring tissue to cause systemic disease. Going forward we plan to collect additional tissue measurements to confirm the results we found in a single pancreas across diverse patients.

## Acknowledgements

This work was funded by the NIH HubMAP initiative grants NIH UH3CA255132 to J.L., and NIH U54DK127823 to M.C.T. and C.M. Mass spectrometry was performed in the Environmental Molecular Sciences Laboratory, a U. S. Department of Energy Office of Biological and Environmental Research national scientific user facility located at Pacific Northwest National Laboratory in Richland, Washington. Pacific Northwest National Laboratory is operated by Battelle for the U.S. Department of Energy under Contract No. DE-AC05-76RLO 1830.

## Data Availability

Raw data has been uploaded to massive and is available at https://massive.ucsd.edu/ProteoSAFe/dataset.jsp?task=a49b0c275ed64142b3c158231bb505ef. Processed data is available at http://synapse.org/microPotsPanc. A Synapse account is required to access the data. All code to analyze the data and recreate the analysis is available at http://qithub.com/pnnl-compbio/hubmap.

---

**Figure S1:**
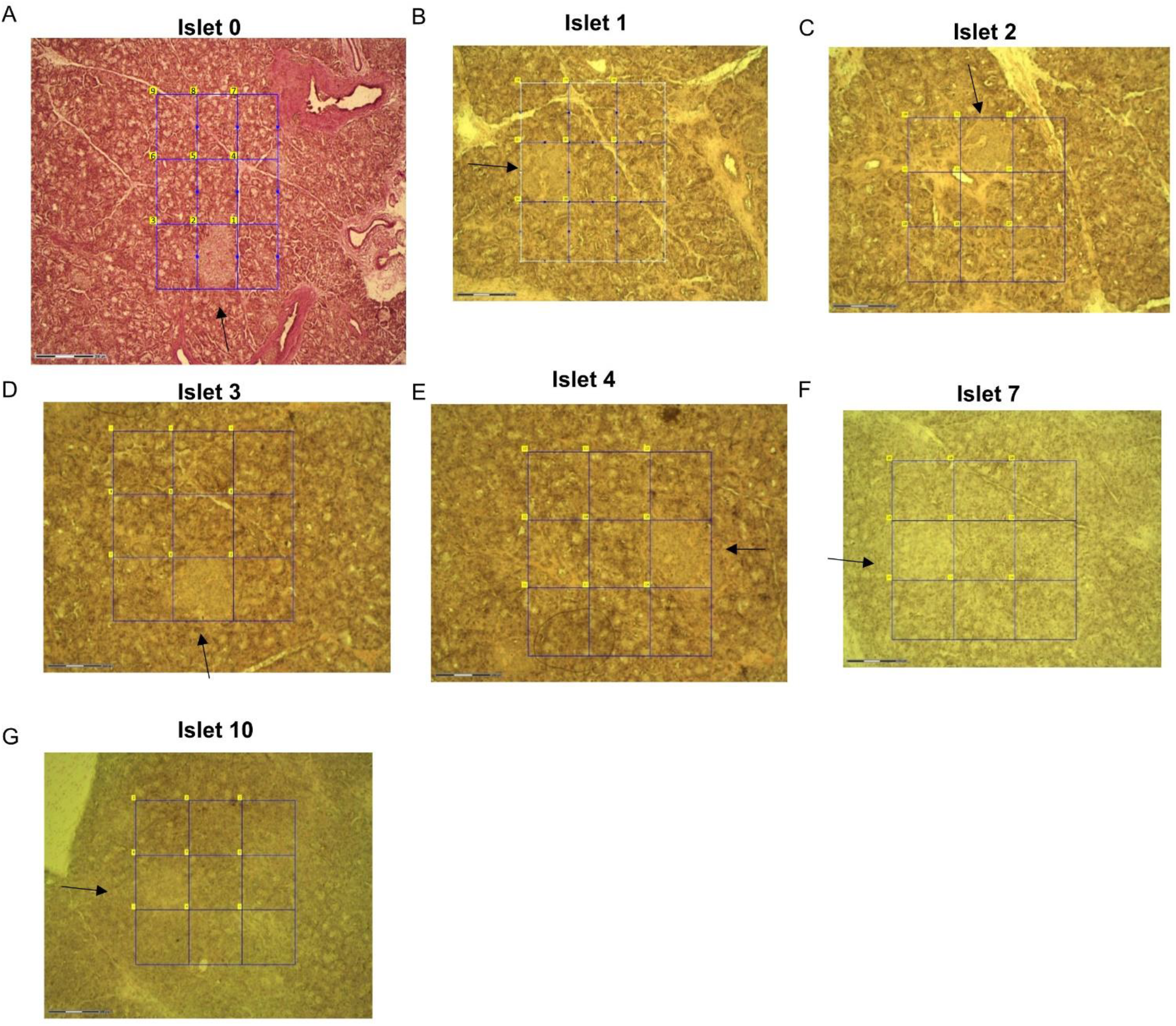
The title of a supplementary figure. Description of pancreas sampling and 3×3 grid. (A) image and islet position of image 0. (B) image and islet position of image 1. (C) image and islet position of image 2 (D) image and islet position of image 3. (E) image and islet position of image 4. (F) image and islet position of image 7. (G) image and islet position of image 10.

